# High-Dimensional Multi-Omic Mapping of Post-Mortem Human Brain Using Iterative Indirect Immunofluorescence Imaging on Xenium-Processed Tissues

**DOI:** 10.64898/2026.09.16.751033

**Authors:** Berke Karaahmet, Tsering Lama, Anqi Wang, Wenqing Cao, Vilas Menon, Hans-Ulrich Klein, David A. Bennett, Philip L. De Jager, Ya Zhang, Mariko Taga

## Abstract

Spatial transcriptomics approaches provide crucial insights into gene expression distribution within intact tissue architecture, but they encounter limitations in detecting morphologically complex cell types, assessing their spatial associations with pathology, and accurately annotating cell types using RNA data alone. Therefore, we developed a robust post-processing workflow integrating Xenium spatial technology with iterative indirect immunofluorescence imaging (4i) on formalin-fixed paraffin-embedded (FFPE) human brain tissue. The post-Xenium 4i protocol presented here allows multi-omic tissue mapping, enabling deeper investigation of pathological microenvironments defined by the spatial distribution of neuropathological hallmarks and enrichment of specific cell populations.

We applied this workflow on calcarine cortex tissue sections where cerebral amyloid angiopathy (CAA) burden is present, in addition to amyloid plaques and tau pathology, generating a 15-plex image that captures the complexity of the pathological microenvironment.

**MOTIVATION:** The expansion of spatial transcriptomics (ST) technologies has enhanced our ability to dissect the complexity and diversity of the molecular organization of the human brain. Among these techniques, the Xenium platform enables high-plex, *in situ* gene expression profiling at subcellular resolution while preserving tissue architecture, thereby offering powerful insights into tissue organization. However, transcriptomic data can be greatly enriched by the inclusion of morphological, anatomical, pathological, and cellular markers at the protein level.

Here, we report a post-spatial profiling workflow that combines the Xenium platform with iterative indirect immunofluorescence imaging (4i) on formalin-fixed paraffin-embedded (FFPE) human brain tissue to capture key pathological features for Alzheimer’s disease (AD) and facilitate cell segmentation. This approach enables cost-effective and flexible multiplexed, multi-omic mapping of the same tissue section by registering Xenium transcript data with subsequent 4i immunostaining, which maximizes the RNA quality. Incorporating immunofluorescence (IF)-based cell-type markers alongside RNA profiles enables cell types to be annotated using robust protein markers rather than relying solely on transcriptomic information, thus reducing an important source of noise in downstream analyses. Immunofluorescence staining of FFPE tissue following Xenium processing also preserves cellular morphology, allowing the morphological complexity of distinct cell types to be captured and quantitated, especially for glial cells with highly ramified processes. To demonstrate the practical application of this workflow to AD studies, we developed a proof-of-concept deep learning framework to classify nuclei into specific major cell types of the brain using the IF data. Thus, we present an integrated experimental and analytic workflow optimized for the multi-omic characterization of human brain tissue in an AD context; these pipelines provide spatially resolved insights into pathological microenvironments found in the older human brain and can readily be adjusted to detect other proteins of interest or other pathological features. Altogether, this approach provides a comprehensive view of spatial architecture within pathologically affected regions and enables accurate transcript-independent cell-type annotation.

**Highlights:**

- Integrate Xenium and 4i to build a multi-omic map on the same FFPE brain tissue
- Improvement of cell-type annotation
- Identify pathologic features and cross-register them into the transcriptomic and proteomic spatial matrix

## INTRODUCTION

Recent single-cell (scRNA-seq) and single-nucleus RNA-sequencing (snRNA-seq) studies have identified distinct cell subtypes associated with disease phenotypes, including those observed in Alzheimer’s disease (AD)^1^. In parallel, recent advances in spatial transcriptomic techniques have addressed a major limitation of scRNA-seq and snRNA-seq: the loss of spatial information, including the spatial relationships between molecular signatures, cell types, and pathologies. Spatial registration enables detailed mapping of specific cell types and their molecular signatures within intact tissue architecture based on transcriptomic profiles. Notably, studies have demonstrated that certain cell populations are differentially enriched in a context-dependent and region-specific manner[1][2][3]. Therefore, integrating transcriptomic data with the broader spatial context is essential to understanding how pathological microenvironments influence the molecular phenotypes of certain cell populations, thereby uncovering potential targets for modulating disease progression.

To date, several spatial transcriptomics platforms are available[4][5][6][7][8][9][10], each with distinct advantages and limitations, and some are compatible with both fresh frozen and formalin-fixed paraffin-embedded (FFPE) tissue samples. Increasingly, spatial transcriptomic approaches have been combined with immunofluorescence (IF) imaging to complement gene expression profiles with protein-level information and histopathological features. In AD, this integration is particularly valuable to map transcriptomic perturbations to neuropathological features of the disease, including amyloid-β (Aβ) plaques and phosphorylated tau aggregates. Previous studies have therefore integrated spatial transcriptomics with IF detection of Aβ and tau to characterize cellular responses within plaque-associated microenvironments. However, many of these studies have relied on a limited number of protein markers or staining of adjacent tissue sections, which can constrain the characterization of multiple pathological features and introduce technical noise due to registration of independent images [11][9]. More recently, methods enabling transcriptomic and protein measurements within the same tissue section have begun to address these limitations. For example, post-Xenium IF staining of IBA1 in FFPE human brain tissue has been used to improve microglial segmentation and transcript assignment[12]. Similarly, post-Xenium staining of neuropathological markers have been used to characterize AD pathology in the Caudate nucleus in Fresh-Frozen (FF) tissue[13]. Furthermore, various modifications to pre-existing sequencing-based spatial transcriptomics techniques have been developed to enable IF or multi-plexed IF imaging on the same tissue section (ST-FFPE-mIF)[14]. Despite these advances, high-dimensional protein imaging together with spatial transcriptomics in the same human brain section remains limited. Such integration is particularly important and useful in AD where pathological context or cell morphologies[15] can enrich or aid in the analysis of spatial transcriptomics data. For example, IF can be used to annotate different cell types or histopathology can be used to determine the spatial association of cells with various inflammatory microenvironments.

To capture the complexity of neuropathology in the context of AD, we optimized a robust protocol that incorporates spatial transcriptomics with the Iterative Indirect Immunofluorescence Imaging (4i) technique, a high-dimensional imaging technique that allows for the multiplexed detection of protein markers on a single tissue section through multiple cycles of staining, imaging, and antibody removal (Figure 1). As proof-of-principle, we developed an IF panel to detect Aβ plaques, tau pathology, vascular abnormalities, and various reactive or homeostatic cellular states. We integrated this 4i panel with the Xenium platform which uses a chemistry that preserves protein epitopes. We show that this approach allows for the generation of a multi-omic map by integrating transcriptomic, multi-plexed IF and histopathological data within a single brain tissue section.

**Figure 1.**
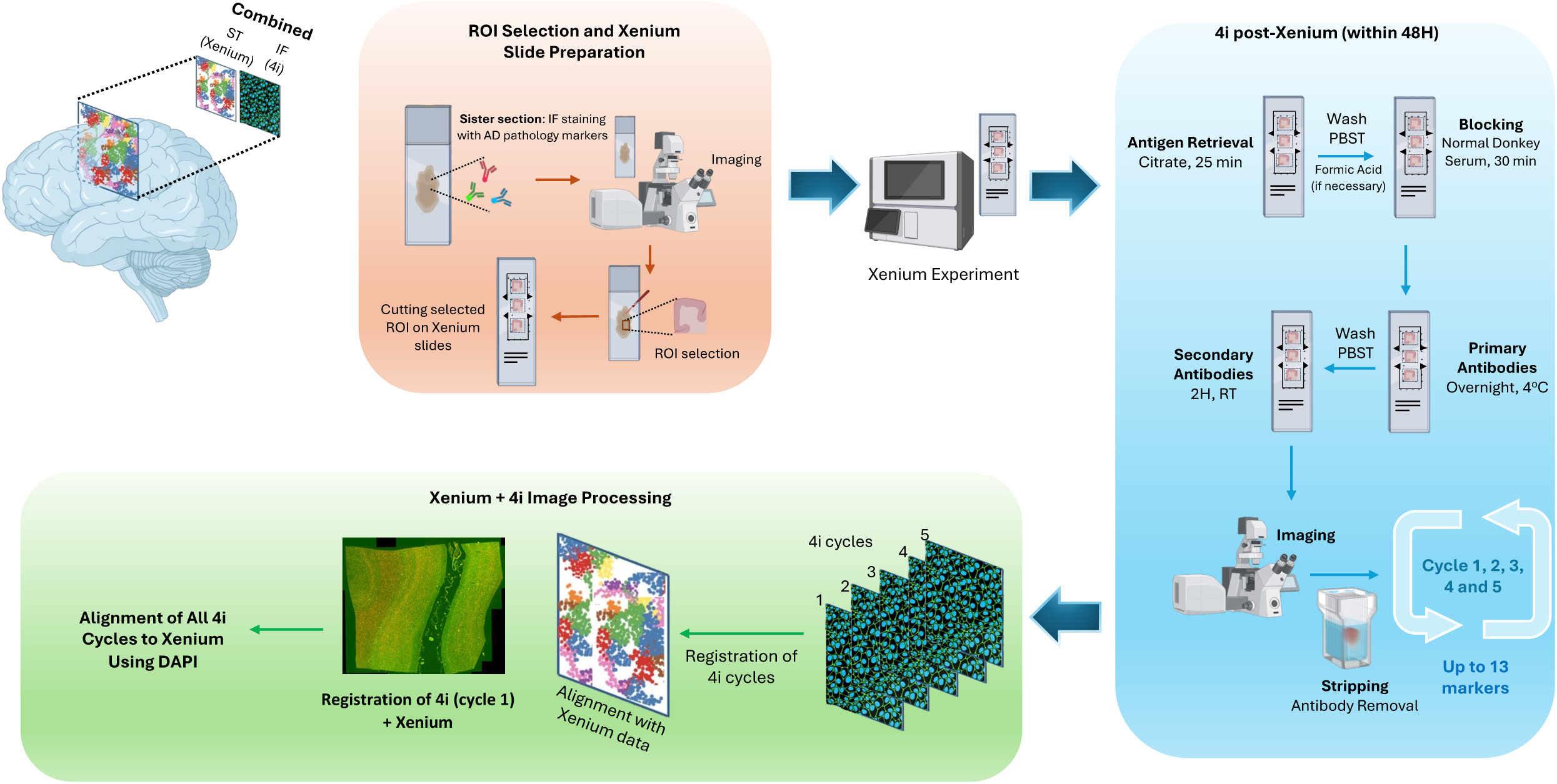
Experimental workflow integrating Xenium with Iterative Indirect Immunofluorescence Imaging (4i) Schematic overview of Xenium combined with the 4i experiment. A sister tissue section was first immunostained for AD pathology markers and images were acquired to select for a ROI. The selected ROI was sectioned and mounted on Xenium slides for the Xenium experiment (up to 3 sections per slide). Within 48H post-Xenium, the same section was processed for 4i, beginning with citrate-based antigen retrieval, followed by blocking, overnight incubation with primary antibodies, and a 2H incubation with secondary antibodies. The entire tissue section was imaged for each cycle. After imaging, antibodies were stripped, and the blocking/staining/imaging workflow was repeated for up to five cycles (up to 20 markers). Downstream image processing included registration of 4i images across cycles and alignment to the Xenium image space using DAPI, enabling integration of spatial transcriptomic and proteomic data.

## RESULTS

### 1. Selection of cellular markers and optimization of the 4i antibody panel

To investigate the spatial microenvironment of pathological processes in neurodegenerative disorders, we developed an integrative method that overlays RNA, pathological features, and protein data within the same tissue section. We designed an antibody panel aimed at identifying the major cell types in the human brain: microglia, astrocytes, excitatory and inhibitory neurons, and oligodendrocytes. Antibodies and their placement across the different 4i cycles were optimized based on the following criteria: (a) high staining specificity, (b) minimal background signal, (c) compatibility with stripping, and (d) stability of staining performance across successive 4i rounds (Supplementary Figure 1). The cellular markers selected for cycle 1 included MAP2 as a pan-neuronal marker, CNP as an oligodendrocyte marker, and NRGN as a cortical excitatory neuronal marker (Figure 2, Supplementary Figure 4). Although all three antibodies were efficiently removed during stripping, their staining qualities were different in later cycles.

**Figure 2.**
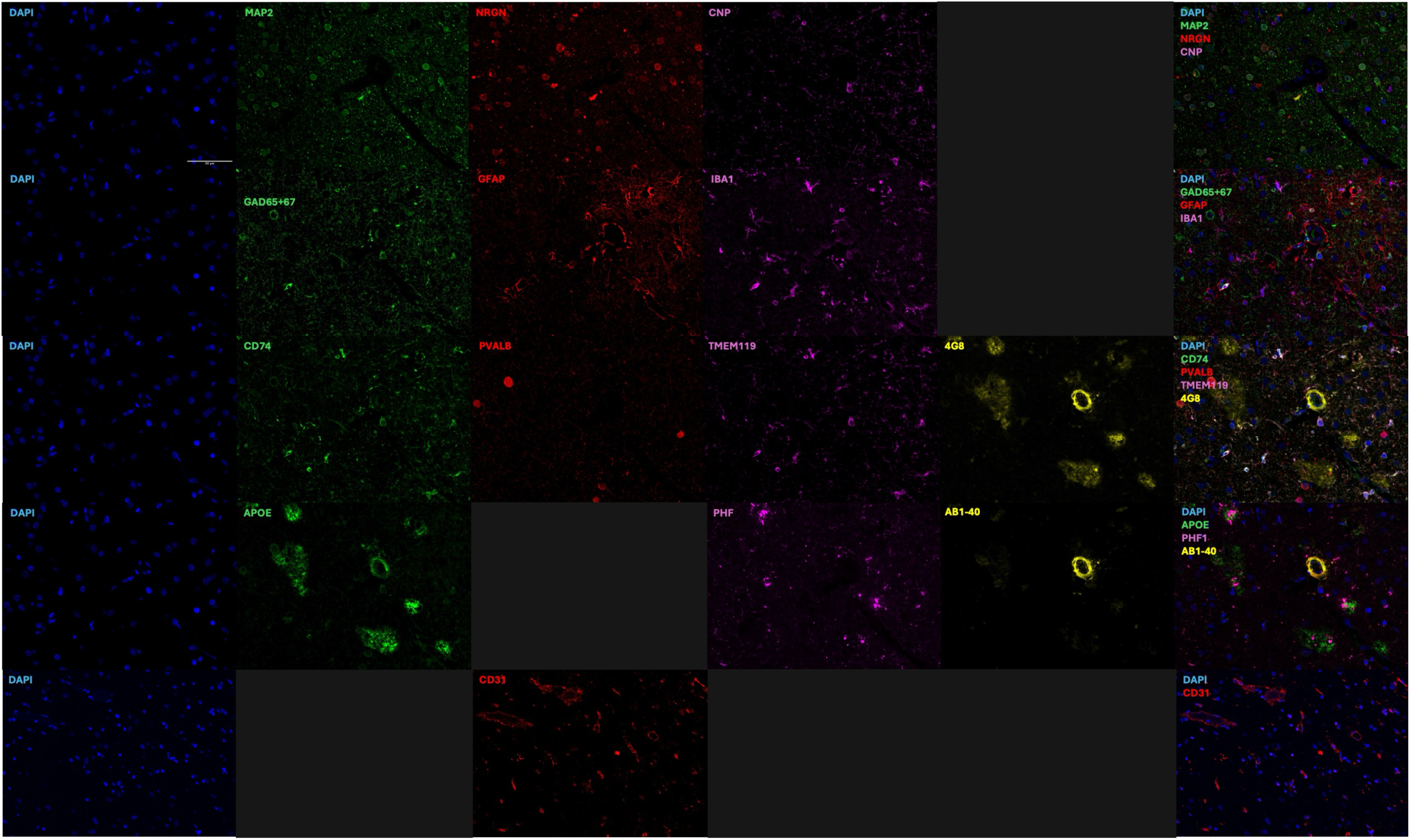
Representative 4i immunofluorescence staining images across 5 staining cycles. Representative immunofluorescence staining images from the calcarine cortex of three individuals with AD pathology. Top to bottom: cycles 1–5, showing single-channel images and the corresponding merged composite (right). Nuclei are labeled with DAPI (blue). Cycle 1: MAP2 (green), NRGN (red), and CNP (magenta). Cycle 2: GAD1/GAD2 (green), GFAP (red), and IBA1 (magenta). Cycle 3: CD74 (green), PVALB (red), TMEM119 (magenta), and 4G8 (yellow). Cycle 4: APOE (green), ptau PHF (magenta), and Aβ_1–40_ (yellow). Cycle 5: CD31 (red). All images were acquired at 20× magnification. All markers were imaged in the same field of view, except for CD31.

MAP2 staining remained stable across cycles, with low background, whereas NRGN and CNP showed relatively reduced specificity and increased background following stripping (Supplementary Figure 1). Therefore, NRGN and CNP were prioritized for cycle 1 of the 4i protocol. For cycle 2, we selected antibodies against GAD1/GAD2 (glutamic acid decarboxylase) to detect cortical inhibitory neurons, along with GFAP for astrocytes, and IBA1 as a microglial/myeloid marker. These antibodies demonstrated efficient removal and maintained high staining specificity following the stripping process (Supplementary Figure 1). Although IBA1 retained good specificity after cycle 2, its staining quality and intensity varied substantially across samples. Similarly, GAD1/GAD2 showed reduced staining performance when tested in later cycles, and, thus, to ensure consistent signal detection, both markers were positioned in the second cycle. For cycle 3, we selected CD74 for MHC^high^ microglia, PVALB (parvalbumin) for one canonical class of inhibitory neurons, TMEM119 (transmembrane protein 119) to distinguish microglia from infiltrating macrophages, and 4G8, to detect Aβ_17-24_ aggregates. All markers retained stable and specific staining performance following two prior stripping rounds (Figure 2). For cycle 4, we selected APOE, PHF for neurofibrillary tau tangles, and Aβ_1-40_, an important pathological marker for CAA. These markers produced high staining quality but, along with 4G8 on cycle 3, showed limited stripping efficiency (Supplementary Figure 2). Therefore, markers of neuropathology were preferentially placed at the end of the staining sequence. For cycle 5, we selected CD31 (cluster of differentiation 31) to label endothelial cells and mark the tissue vasculature. CD31 maintained robust staining after four prior staining and stripping rounds. However, PHF, stained in the previous round, was raised in the same host species and exhibited insufficient stripping, resulting in residual cross-cycle reactivity. Our empirical testing demonstrated that antibody performance and overall 4i assay robustness were strongly influenced by cycle order, highlighting the importance of strategic antibody panel design.

A β-mercaptoethanol (βME)-based stripping buffer enables efficient antibody elution while preserving antigenicity and tissue integrity across multiple staining cycles. When compared with a glycine-based stripping buffer, βME consistently removed antibodies, preserved nuclear DAPI signal and showed minimal residual signal from previous staining rounds, indicating a more effective and broadly applicable stripping buffer (Supplementary Figure 3). The βME stripping buffer, adapted from Cattoretti et al. (2004)[16], was optimized for the post-Xenium 4i workflow to support reproducible multiplexed detection for our custom antibody panel. Given the lipid-rich nature of FFPE human brain tissue and susceptibility to epitope masking, we modified the stripping conditions by increasing the incubation temperature to 60°C and extending the incubation time to 45 minutes.

### 2. Sample Quality Control and Region-Of-Interest selection

Before the Xenium and 4i experiments, adjacent FFPE tissue sections underwent standardized pre-assay sample Quality Control (QC) and region-of-interest (ROI) selection. These sections were deparaffinized and then subjected to histological and molecular quality checks, including hematoxylin and eosin (H&E) staining to assess morphology, DAPI staining to evaluate nuclear integrity, and immunofluorescence staining to guide ROI selection to enrich for neuropathological features. These QC steps enabled selection of regions with preserved morphology and relevant protein marker expression, ensuring that chosen ROIs were biologically meaningful and suitable for high-quality Xenium and 4i analyses (**Materials and Methods**). The ROI was defined by the enrichment of amyloid plaques (including both diffuse and neuritic plaques) and CAA in the adjacent tissue section. We ensured that each ROI encompassed the six cortical layers and a portion of the underlying white matter. All selected samples were stained with Aβ_17–24_, which detects the pathologies relevant to this study. Because we aimed to fit three tissue sections on a single Xenium slide (video at ref 19), section size was constrained; however, each section was large enough to include neuritic plaques, diffuse plaques, parenchymal CAA, and meningeal CAA, whenever these features were present. For samples without AD pathology, we preferentially selected a region containing both meningeal and parenchymal blood vessels so that they could serve as non-pathological controls for comparisons with AD pathology samples.

### 3. Immunohistochemistry and 4i staining post-Xenium

All Xenium slides are recommended to undergo immunohistochemistry (IHC) within 48 hours and be stored in PBS-T at 4°C. For longer storage up to one week, PBS-T should be replaced regularly every 3 days. Here, we performed 4i using heat-induced antigen retrieval step with citrate buffer (pH 6.0; 25 min at 400 W). In these experiments, antigen retrieval did not compromise staining quality for any of the markers. If background reduction is a priority, the step can be removed, but only after confirming that all antibodies in the panel perform adequately without antigen retrieval. We also did not observe any tissue detachment from the slides following the antigen retrieval step. In addition, other pre-treatments, such as formic acid required for Aβ staining, can be introduced before any cycle in 4i. Following a full cycle of staining, the tissue sections were scanned, and the slides underwent BME–based antibody stripping **(Materials and Methods)**. After each stripping step, antibody removal was verified by microscopy, followed by a blocking step to prepare for the next staining cycle (Figure 2). The coverslip was removed gently at each step to avoid tissue damage or microtears.

### 4. 4i staining post-Xenium whole tissue imaging

After each staining cycle, the entire section was imaged using an immunofluorescence microscope. The same imaging pipeline was applied across all sections and all 4i cycles to ensure consistency. A semi-automated acquisition pipeline (JOBS) was created in NIS-Elements AR for high-resolution, high-throughput imaging of whole tissue sections. Autofocus at 4x BF initiated a prescan over the entire slide to generate a low-resolution overview map for region of interest (ROI) selection. To scan the ROI, the region was manually annotated and autofocus was performed over a 600 µm continuous scan at 20x, followed by a 1 µm offset deeper into the tissue for fine-tuning. The autofocus algorithm excluded the outer 20% of the tissue border to avoid edge artifacts.

To make use of Extended Depth of Focus (EDF) algorithm, z-stacks were captured at each imaging position. Four z-stacks separated by 1.5 µm per plane were acquired. This approach accounted for grooves, folds, and sectioning artifacts that caused uneven tissue surfaces. All images were saved in Nikon Digital version 2 (.nd2) format, and individual fields of view were stitched and converted to .ome.tif format using custom scripts for broad accessibility **(Materials and Methods)**.

### 5. Application of deep learning to identify major cell types Xenium-detected nuclei

Accurate cell-type classification in spatial transcriptomics often relies on RNA-based *de novo* annotation or cross-modality label transfer, both of which can be noisy. Inclusion of cell type specific markers in the 4i panel provides a unique opportunity for classification of each nucleus to the broad cell types of interest, while reserving RNA-based classification for finer-grained phenotypic resolution. To this end, we trained a deep learning classifier using human-annotated nuclei from 15 individual xenium datasets (Figure 3a, Supplementary Figure 5a, b, Supplementary Table 1). As proof-of-principle, we hypothesized that a classifier trained on a subset of nuclei from these Xenium sections could be robustly applied to all the nuclei within the same 15 sections. We chose this approach because deep neural network models have the capability to classify nuclei using multiple markers and morphology simultaneously. To reduce biological and staining heterogeneity, we restricted annotations and downstream analyses to grey matter. The classification categories were Neurons (neu; MAP2^+^), Astrocytes (ast, GFAP^+^), Microglia and Macrophages (mic, IBA1^+^), Oligodendrocytes (oli, CNP^+^), nuclei of vasculature (vas, defined by morphology and spatial context), and unknown (unk), an unclear class in which nuclei do not express any of the 4i markers or have ambiguous signals. Using visual inspection and manual annotation, we generated a training dataset of 80,500 annotated nuclei. 10,909 of these nuceli were held out for validation, and an independent set of 10,492 nuclei were held out for testing the model (Supplementary Figure 5a).

**Figure 3.**
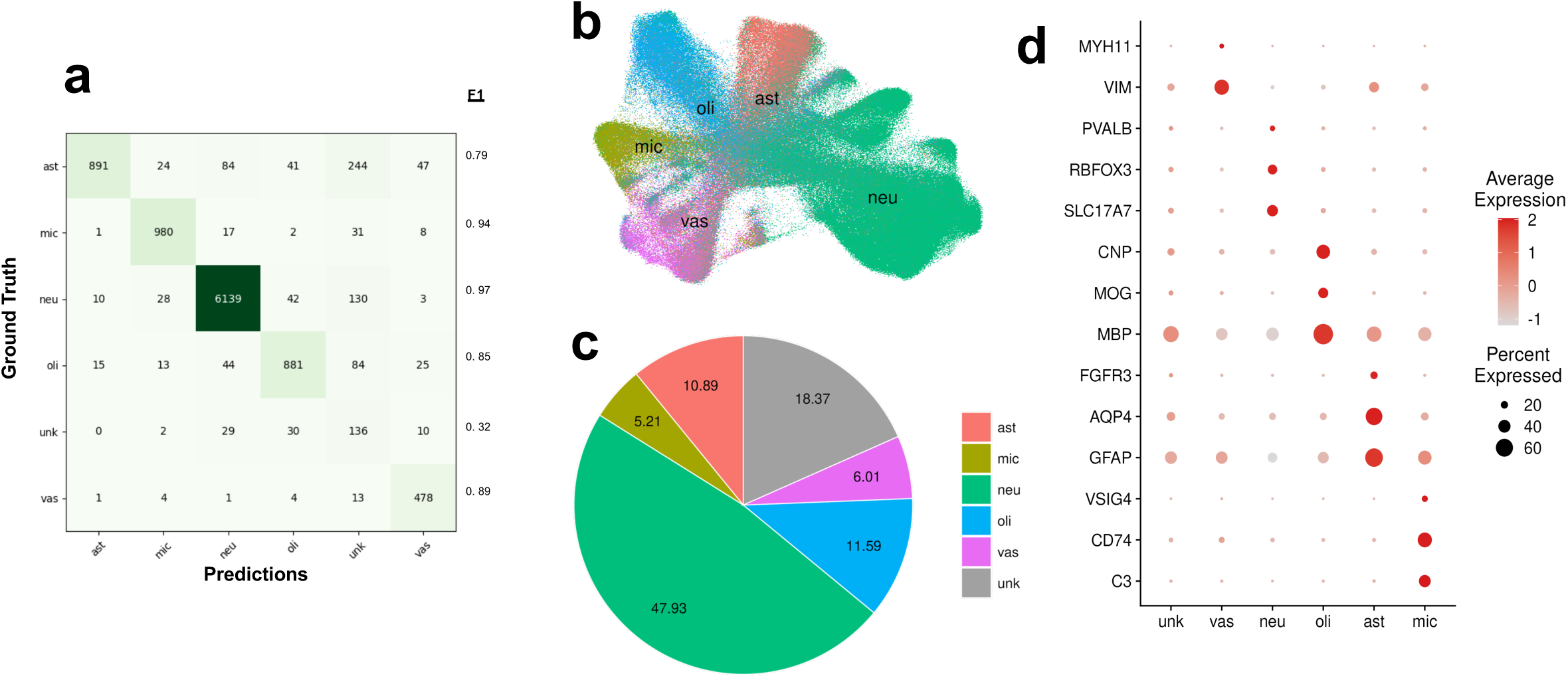
Deep learning-based classification of Xenium-detected nuclei into major cell types of the brain using 4i immunofluorescence reveals distinct RNA profiles. **(a)** Confusion matrix between human labeled annotations of 10,492 cells set aside for testing and deep learning-based classification. **(b)** Proportion of cell types among all nuclei from 15 tissue sections presented in this study. **(c)** UMAP of RNA profiles overlaid with deep learning classifications shown in different colors. Overall, each of the major CNS cell-types cluster into distinct spaces on RNA-based UMAP coordinates. **(d)** Dot plot showing that cell type-specific markers are highly expressed by the respective cell type identified through deep-learning.

After training, we tested our classifier on the test-split (n = 2 samples 10,492 cells). F1-score, which is a common performance metric used in classification tasks with class imbalances, was F1_neu_ = 0.97, F1_ast_ = 0.79, F1_mic_ = 0.94, F1_oli_ = 0.85, F1_vas_ = 0.89, and F1_unknown_ = 0.32 (Figure 3a). These values show a strong predictive power on major cell types and is expected for the unknown class as it acts as a general unknown/unclear category.

We applied this version of the classifier to all of the 538,318 nuclei (DAPI+ objects) in grey matter identified in the 15 xenium sections. RNA-based UMAP projections showed that nuclei predicted to belong to the same cell type by the deep learning classifier were co-localized in the reduced dimensional space (Figure 3b). Out of all the nuclei that were not in the unknown category (81.63% all nuclei), we observed that 58.71% of nuclei were classified as neurons, 6.38% as microglia, 13.35% as astrocytes, 7.36% as vasculature, and 14.20% as oligodendrocytes (Figure 3c). These distributions are consistent with previously reported cellular compositions of human cortical grey matter[17]. Lastly, we confirmed that the RNA expression of cell type specific markers indeed map to our classifier-predicted annotations (Figure 3d).

Information on exact cell shape outlines also provides a unique opportunity to test the quality of cell segmentation as it pertains to assigning specific transcripts. While previous studies have compared various computational approaches for this purpose, they did not have access to ground truth information that combination with 4i data provides[18]. Therefore, by assessing the specificity and sensitivity of a subset of cell-type specific transcripts, we assessed the transcript classification performance of 5 different approaches to generating a cell mask: manual marker-guided segmentation, nucleus-only segmentation, 5µm expansion of the nucleus and 50% or 100% expansions of the nucleus (**Materials and Methods**). As expected, we observed that nucleus-only segmentation captured the least number of transcripts, regardless of specificity, compared to other segmentation methods. Importantly, we found that the 5µm approach performs very similarly to the manual marker-guided segmentation mask for microglia (F1_manual_ = 0.53, F1_5µm_ = 0.54), oligodendrocytes (F1_manual_ = 0.47, F1_5µm_ = 0.42), astrocytes (F1_manual_ = 0.30, F1_5µm_ = 0.33) and neurons (F1_manual_ = 0.69, F1_5µm_ = 0.77). Furthermore, the 5µm expansion approach performed better than smaller expansions of the nucleus (i.e. 50% expansion) or nucleus segmentation alone (Supplementary Figure 6a-b). While the interpretation of these results heavily depends on cell types, protein markers used to define the boundaries, and transcripts of interest; 5µm expansion of nucleus balances false positives and negatives as effectively as exact cell segmentation for the scope of this study.Taken together, these results demonstrate an extension of the standard Xenium workflow through integration of a 4i panel in human FFPE brain tissue, enabling robust cell-type classification of nuclei using histological markers through deep learning and identification of histopathological features.

## DISCUSSION

Advances in spatial technologies now enable detailed investigation of the cellular composition of the pathological microenvironment and characterization of the spatial relationships between these cell populations and pathological features. However, limitations remain when integrating pathological markers and proteins of interest, because RNA abundance does not always reflect protein levels. Challenges also arise when incorporating cellular morphology into analyses, since some cell types undergo activation-associated morphological changes. In addition, it is often necessary to include multiple markers that cannot be accommodated within a single round of staining. Here, we establish a robust workflow covering the post-Xenium immunofluorescence with multiple rounds of staining and cell-type annotation using immunofluorescence information, addressing challenges commonly encountered with current approaches.

While standard immunofluorescence staining protocols can be integrated into the workflow described here, a key consideration is to initiate immunofluorescence staining within 48 hours. If necessary, slides can be stored for up to one week by keeping them immersed in PBS-T and replacing the buffer every three days. Determining the antibody staining order is also a key step in this study, enabling the inclusion of key cellular markers that were subsequently used for cell annotation and neuropathological examination. However, we do not recommend using immunofluorescence intensity measurements from later staining cycles to address biological questions, as repeated stripping may affect antibody binding and increase tissue background.

A variety of antibody elution buffers have been reported in the literature. In this study, we tested two stripping protocols: one based on BME and the other based on a HCl-glycine solution. We selected the BME-based protocol because it allowed for additional staining cycles while maintaining low background and specific signal detection, including IBA1 and DAPI (Supplementary Figure 3). The selection of cell-type markers was challenging because we prioritized cytoplasmic markers, particularly for glial cells. Since IBA1 is not entirely specific to microglia, TMEM119 and CD74 can be included to improve microglial specificity. In addition, markers such as CD74 carry biological significance because of their reported involvement in AD pathology and cognitive impairment[19][20]. Although ALDH1L1 is widely used as a general astrocyte marker, we prioritized GFAP for astrocyte detection because its expression intensity varies with astrocyte reactivity. The selection of neuronal markers also required careful consideration. We selected MAP2 over NeuN because MAP2 provides stronger cytoplasmic staining, whereas NeuN primarily labels the nucleus. However, we also acknowledge that MAP2 signal may be reduced in a subset of phospho-tau–positive neurons, particularly in pathological brains, where phospho-tau accumulation is associated with dendritic and somatic structural disruption[21]. Therefore, inclusion of additional neuronal markers may further improve the detection of neuronal populations, if these markers can be incorporated into the 4i panel.

This workflow also addresses current limitations in cell-type annotation, enabling us to move beyond regional analyses toward more cell-specific analyses. To support this, we developed a proof-of-concept method for classifying Xenium-detected nuclei into major central nervous system cell types using 4i-derived data. Because RNA quality can vary depending on tissue quality and preservation method, including whether samples are FFPE or fresh frozen, or simply due to limited sizes of panels, this approach may be particularly useful for identifying and annotating broad cell types.

Deep learning models can accommodate moderate inter-sample staining variability while leveraging additional features beyond marker intensity, including cellular morphology and spatial context. Therefore, using 4i-derived immunofluorescence data for cell-type classification may reduce the need to allocate space in the Xenium panel to broad cell-type markers. Given the 480-gene limit of our panel, this approach can allow RNA targets to be prioritized for disease-associated genes and cell-subtype-specific markers. Additionally, the use of immunofluorescence for identification of broad cell types is invariant to RNA quality, which can be an advantage in instances with low tissue quality or FFPE preservation.

An alternative use of this multi-modal data could be to extract accurate cell segmentation masks. While this can increase accuracy of transcript assignments, it could also introduce additional noise due to overlapping cellular processes. In our experiments, we observed most of the RNA molecules to be inside or immediately proximal to the nucleus. Therefore, using nucleus-only or slightly expanded nucleus masks, combined with immunofluorescence-based cell type annotation, can generate data that is ready for downstream differential expression analysis.

In conclusion, this study establishes a robust and scalable workflow for integrating post-Xenium 4i with spatial transcriptomics data in human post-mortem FFPE brain tissue. By optimizing the immunofluorescence staining workflow, including antibody order, antibody elution, and downstream image analysis, this approach enables the simultaneous assessment of cell-type identity, neuropathological features, proteins of interest and transcriptomics within the same tissue section. Importantly, the integration of 4i-derived protein data can be used for cell-type annotation, particularly when classification based on transcriptomic data alone is limited. Together, this spatial method and workflow provide a powerful framework for deeper investigation of pathological microenvironments. By enabling the identification of target genes associated with distinct pathological features, this approach can help elucidate mechanisms involved in pathology formation and support the characterization of pathology subtypes and their differences.

## FIGURE LEGENDS

**Supplementary Figure 1. Representative 4i immunofluorescence staining images of all markers across 5 staining cycles.**

Representative immunofluorescence staining images from the dorsolateral prefrontal cortex of an individual with AD pathology. Top to bottom: all cell-type markers in 4i antibody panel, showing assigned single-channel images of each marker stained across five cycles. Markers in descending order: MAP2, (green), CNP (magenta), GFAP (red), IBA1 (magenta), NRGN (red), CD74 (green), TMEM119 (magenta), PVALB (red), APOE (green), CD31 (red), GAD1/GAD2 (green). All images were acquired at 20× magnification.

**Supplementary Figure 2. Representative 4i immunofluorescence images showing 4G8 and PHF signals after stripping.** Fluorescence remaining after treatment with the β-mercaptoethanol-based stripping solution indicated incomplete antibody removal. Therefore, 4G8 and PHF were assigned to later 4i cycles to minimize interference with antibody staining in subsequent rounds. All images were acquired at 20× magnification.

**Supplementary Figure 3. Representative 4i immunofluorescence staining images of DAPI and IBA1 in β-mercaptoethanol vs glycine stripping solution across 5 staining cycles.**

Representative immunofluorescence staining images from the dorsolateral prefrontal cortex of an individual with AD pathology. Nuclei are labeled with DAPI (blue), and microglia are labeled with IBA1 (magenta). **(a)** β-mercaptoethanol stripping solution used for antibody elution between cycles. **(b)** Glycine stripping solution was used for antibody elution between cycles. All images were acquired at 20× magnification. All markers were imaged in the same ROI.

**Supplementary Figure 4. Additional representative 4i immunofluorescence images across 5 staining cycles.** Representative immunofluorescence staining images from the calcarine cortex of three additional individuals with AD pathology, demonstrating staining consistency across donors. Top to bottom: cycles 1–5, showing single-channel images and the corresponding merged composite (right). Nuclei are labeled with DAPI (blue). Cycle 1: MAP2 (green), NRGN (red), and CNP (magenta). Cycle 2: GAD1/GAD2 (green), GFAP (red), and IBA1 (magenta). Cycle 3: CD74 (green), PVALB (red), TMEM119 (magenta), and 4G8 (yellow). Cycle 4: APOE (green), ptau PHF (magenta), and Aβ_1–40_ (yellow). Cycle 5: CD31 (red). All images were acquired at 20× magnification. All markers were imaged in the same ROI, except for CD31.

**Supplementary Figure 5. Training of the deep learning classifier. (a)** Numbers of manually annotated nuclei belonging to each class used for training and validation from 15 tissue sections. **(b)** Training metrics showing loss and accuracy over epochs.

**Supplementary Figure 6. Assessment of different segmentation masks on transcript assignments. (a)** Representative images showing an example of a given cell, nucleus based segmentation masks (bubbles: 5µm expansion, outer contour: 100% expansion, inner contour: 50% expansion, innermost contour: nucleus only segmentation), manual segmentation outlined by a human observer, transcripts specific to that cell type and transcripts specific to other cell types. **(b)** Quantification of average F1-score across the samples. n = 3 tissue sections. Genes: n_mic_ = 32, n_oli_ = 7, n_ast_ = 6, n_neu_ = 19, n_vasc_ = 4. Cells: n_mic_ = 40-41, n_oli_ = 18-25, n_ast_ = 15-30, n_neu_ = 40-45 per tissue. Scale bar 10μm.

**Supplementary Table 1. Demographics**

Post-mortem calcarine cortex brain tissue from 15 donors was used in this study. The table lists the subject ID, age at death (years), clinical AD diagnosis, neuropathological AD diagnosis, CAA severity (mild/moderate/severe), and sex.

**Supplementary Table 2. 4i antibody panel and fluorophore assignments across different staining cycles.**

Table shows the 4i staining design across five cycles, with markers assigned to Alexa Fluor 488, 568, 647, and 790 channels. Cycle 1: MAP2 (AF488), NRGN (AF568), CNP (AF647). Cycle 2: GAD1/GAD2 (AF488), GFAP (AF568), IBA1 (AF647). Cycle 3: CD74 (AF488), PVALB (AF568), TMEM119 (AF647), β-amyloid 17–24 (AF790). Cycle 4: APOE (AF488), phospho-tau S396 (AF647), β-amyloid 1–40 (AF790). Cycle 5: CD31 (AF568).

**Supplementary Table 3. List of antibodies used for 4i staining, including dilution factors, catalog numbers, and manufacturers.** Antibody dilutions were optimized for the specific staining cycle in which each antibody was applied.

## MATERIALS AVAILABILITY

This study did not generate new unique reagents.

## DATA AND CODE AVAILABILITY

All the code used in this study can be accessed at: https://github.com/cu-ctcn/cascade/tree/main/xenium4iMethodsScripts. Xenium and post-Xenium 4i data can be accessed at https://www.synapse.org/Synapse:syn68493629 upon request.

## MATERIALS AND METHODS

### ROSMAP cohort

The Religious Orders Study (ROS) and Rush Memory and Aging Project (MAP) are both cohort studies conducted by the Rush Alzheimer’s Disease Center in Chicago. Older individuals free of known dementia at enrollment agreed to annual clinical evaluation and brain donation at the time of death. Both studies were approved by an Institutional Review Board of Rush University Medical Center[22]. All participants signed informed and repository consents and an Anatomic Gift Act.

Both cohorts have a large common core of identical data at the item level, including 21 cognitive functional tests, using validated procedures to diagnose AD and other dementias. All participants undergo a standardized structured assessment for AD which includes CERAD, Braak Stage, NIA-Reagan, and a global measure of AD pathologic burden with modified Bielschowsky. β−amyloid load was measured by staining three monoclonal antibodies against Aβ, which are 4G8, 6F/3D and 10D5, while PHF Tau tangle density was measured using the AT8 antibody across eight brain regions. Both frozen and fixed brain tissues are accessible upon request from these subjects. AD pathology was assessed according to the National Institute on Aging-Reagan criteria. The clinical and pathologic methods have been previously reported[22][23].

### QC and ROI selection

Tissue sections (5 µm thick) were incubated at 60°C for 30 min, then cooled at room temperature (RT) for 7 min. Sections were deparaffinized in xylene for 10 min twice, followed by incubation in 100% ethanol for 3 min twice and 96% ethanol for 3 min twice. Sections were then incubated in 70% ethanol for 3 min, washed with Milli-Q water for 20s, and immediately processed for either immunofluorescence or H&E (Hematoxylin and Eosin) staining. For the H&E staining, sections were incubated in eosin solution (a 400:1 mixture of eosin and 0.1 M hydrochloric acid) for 3 min followed by incubation in 95% ethanol for 30s twice, then in 100% ethanol for 30s twice, and cleared in xylene for 3 min twice. Slides were allowed to dry completely and mounted by adding ∼100 µL ProLong Glass onto the tissue section. For immunofluorescence-based ROI selection, tissue sections were washed three times with 1× phosphate-buffered saline (PBS) at RT. Heat-induced antigen retrieval was then performed in citrate buffer using a microwave (800 W, 30% power) for 25 min, followed by three washes in 1× PBS at RT. Sections were blocked with 3% bovine serum albumin (BSA) for 30 min at RT and incubated overnight at 4°C with primary antibodies diluted in 1% BSA in 1× PBS. The next day, sections were washed three times with 1× PBS at RT and incubated with Alexa Fluor–conjugated secondary antibodies diluted in 1% BSA in 1× PBS for 1H at RT. After three additional washes in 1× PBS at RT, slides were mounted with ∼100 µL of an anti-fade mounting medium containing DAPI. The selected ROI was approximately 8 mm × 6 mm to accommodate three sections within the capture area.[24]

### Xenium in situ workflow

#### Gene panel design

A custom 480-gene probe panel was designed for cell type identification, based on the Xenium human brain panel and in-house scRNA-seq data from human brain tissue. The panel targets major brain cell types – including neurons, microglia, oligodendrocytes, and astrocytes – and includes negative controls. Each probe consists of two complementary sequences that hybridize to the target RNA and a third region encoding a gene-specific barcode. Upon hybridization, the probe ends ligate to form a circular DNA molecule. This design ensures high specificity: off-target binding events are suppressed because ligation requires perfect complementarity, thereby minimizing background signal.

#### Xenium sample preparation

Formalin-fixed, paraffin-embedded (FFPE) tissue blocks were sectioned at 5 µm and mounted onto Xenium slides (PN-1000465) following an established protocol[25][24]. Slides were air-dried at RT for 30 minutes, then incubated at 42 °C for 3 hours on a Xenium Thermocycler Adapter plate. After overnight storage with desiccant at RT, slides were processed using the Xenium In Situ for FFPE – Deparaffinization and Decrosslinking protocol (10X Genomics, CG000580 Rev C). Slides were assembled into Xenium cassettes (PN-1000566) to enable precise temperature control in a PCR instrument. Using the Xenium Slides and Sample Prep Reagents kit (PN-1000460), slides were incubated in decrosslinking/permeabilization solution at 80 °C for 30 minutes. Subsequent steps followed by the Xenium In Situ Gene Expression user guide (CG000582 Rev D). Hybridization was performed at 50 °C for 17 hours with the custom 480-gene panel. Post-hybridization washes and enzymatic steps included a 30-minute wash at 37 °C, 2-hour ligation at 37 °C, and 2-hour amplification at 30 °C. Finally, slides were washed, treated with an autofluorescence quencher, and stained with DAPI.

#### Xenium Analyzer instrument

Processed slides in Xenium cassettes were imaged using the Xenium Analyzer according to the Xenium Analyzer User Guide (CG000584 Rev B). Two slides were loaded per run, and an initial sample scan generated low-resolution images of fluorescent nuclei. The user then manually selected regions of interest for high-resolution analysis.

The Xenium Analyzer is a fully automated instrument that decodes RNA targets at subcellular resolution. After loading slides and consumables, the system controls all fluid handling and experimental steps. Data collection proceeds through repeated cycles of fluorescent probe binding, image acquisition, and probe stripping. Onboard analysis pipelines (version 1.8.2.1, software version 1.7.1.0) assign confidence scores to each detected transcript. Output images and expression profiles were evaluated using Xenium Explorer software (version 2.0.0).

### Post-Xenium sample processing

Iterative indirect immunofluorescence staining was performed on 5μm post-Xenium sections of formalin-fixed paraffin-embedded (FFPE) human brain tissue from the calcarine cortex. Post-Xenium slides with FFPE tissue sections can be stored in PBS containing Tween 20 (PBS-T) at 4°C for up to 7 days. Tissue sections were washed with PBS-T before immunofluorescence staining.

#### Iterative indirect immunofluorescence staining

Heat-induced epitope retrieval was performed using citrate buffer (cat#C9999, Millipore Sigma) in a microwave (1200W, 30% power) for 25 minutes. The tissue sections were then washed with PBS-T and blocked with 5% normal donkey serum (NDS, cat#017-000-121, Jackson ImmunoResearch) for 30 minutes at RT. When a primary anti-β-Amyloid was used, the tissue was incubated in 88% formic acid for 2 minutes prior to blocking.

For immunofluorescence staining, tissue sections were incubated overnight at 4°C with primary antibodies diluted in 1% NDS prepared in 1X PBS (cat#119069131, Quality Biological). The following day, slides were washed with PBS-T and incubated with Alexa Fluor-conjugated secondary antibodies (1:500 dilution in 1% NDS/PBS; Invitrogen) for 1 hour at RT. For detection using conjugated primary antibodies, sections were washed with PBS-T, blocked with 5% NDS for 30 minutes at RT, and then incubated with the conjugated primary antibody diluted in 1% NDS/PBS for 1 hour at RT. Following immunolabeling, slides were treated with 1X TrueBlack lipofuscin quencher (cat#23007, Biotium) for 2 minutes at RT to minimize autofluorescence. Slides were mounted with an anti-fading reagent containing DAPI (cat#P36931, Invitrogen) and coverslipped (cat#CG15KH, ThorLabs Inc.).

After each round of staining and image acquisition, slides were incubated in an antibody stripping buffer for 45 minutes at 60°C. The stripping buffer contained β-mercaptoethanol (βME, cat#63689, Sigma-Aldrich), 10% sodium dodecyl sulfate (SDS) solution (cat#AM9822, Invitrogen), 1 M Tris hydrochloride (Tris-HCl, pH= 6.8, cat#ab286853, Abcam), and Milli-Q water (mqH2O). A 100 mL elution buffer was prepared by adding 20 mL of 10% SDS solution and 6.25 mL of 1 M Tris-HCl (pH= 6.8). Then the volume was brought to 100 mL with 60°C heated mqH2O. The pH of the elution buffer was confirmed to be 6.8 before adding 800 μL of βME. Following incubation in the stripping buffer, slides were washed with mqH2O for one hour at RT on a shaker, with water changes every 15 minutes. The slides were then incubated in tris-buffered saline with Tween (TBS-T, cat#28360, Thermo Scientific) for 5 minutes at RT, followed by a 5-minute treatment with 1× TrueBlack lipofuscin autofluorescence quencher at RT. Slides were mounted using ProLong Glass antifade reagent (cat#P36984, Invitrogen) for confirmation of antibody elution under a fluorescent microscope. The next iteration of immunofluorescence staining was performed after tissue sections were washed with PBS-T and blocked with 5% NDS for 30 minutes at RT. The detailed protocol is described in Lama T. et al., 2026[26].

#### Creation of Nikon JOBS for Post-Xenium Tissue Scanning

The Nikon Eclipse Ni-E epifluorescence microscope performed semi-automated whole-section imaging using NIS-Elements Advanced Research software. Image acquisition was executed through the JOBS module, which enabled a flexible, customizable workflow using a visual programming interface. This drag-and-drop system allowed users to create acquisition pipelines without writing code. Images were stored in ND2 format along with associated metadata output. Using this interface, we designed an automated, high-resolution, high-throughput image acquisition pipeline.

To create the JOBS pipeline, (1) a new project was opened and labeled. (2) Prescan settings were defined by adding a “Capture Definition” task to configure optical settings for 4× Brightfield (BF). (3) A second “Capture Definition” task was added for 20× fluorescent imaging. (4) Exposure times were preset using two “Set Exposure to Optical Configuration” tasks, assigning absolute exposure times of 3 ms and 70 ms for 4× BF and 20× fluorescent channels, respectively. (5) A “Macro” task established camera formatting using the commands ROISet() and CameraFormatSet(). We defined ROISet (252, 252, 2052, 2052) to set the ROI size to 1800 × 1800 pxs. The command “CameraFormatSet(1, “FMT 1×1 16”); CameraFormatSet(2, “FMT 1×1 16”);” configured the camera to 1×1 binning with a 16-bit image format. (6) Autofocus parameters were set for 4× BF and 20× BF optical configurations using two “Autofocus Settings” tasks. Autofocus at 4× BF was performed over a 1500 µm single pass at a slow speed with continuous adjustment, whereas autofocus at 20× BF was performed over a 600 µm single pass under similar conditions. For 20× BF, the z-stack was offset by 1 µm deeper into the tissue for fine-tuning. In both tasks, 20% of the tissue border was excluded to avoid edge artifacts, and the system reverted to the original z position if autofocus failed. (7) A “Get Current XY Position” task saved the starting x-y position in the metadata. (8) Objective and stage positioning were controlled using a “Macro” task. The command “Stg_SetNosepiecePosition(0);” moved the objective to position 0 (4×), and “StgMoveMainZ(1050, 0)” moved the stage to an absolute z-position of 1050 µm. (9) An “Autofocus” task applied the previously defined 4× BF autofocus settings to generate a whole-section overview. (10) A “Scan Large Image” task then acquired and stitched multiple fields of view into a composite image. Using the predefined 4× BF Prescan settings, a region of 40 × 11 tiles was acquired from the top-left position. A 10% overlap and blended stitching mode were applied. (11) A “Draw Regions” task was used to define ROIs on the composite image for subsequent scanning manually. This was useful for the Xenium experimental design, where three tissue sections were mounted within a single capture area. (12) A “Macro” task was used to change the nosepiece position using “Stg_SetNosepiecePosition(2);”, corresponding to the 20× objective. (13) A “Select Optical Configuation” task reset the system to 4× BF following image acquisition. (14) A “Move to XY Position” task returned the stage to the initial x-y position saved earlier. (15) A “Storage” task named and saved the acquired large image to the selected folder.

#### Image Acquisition

Semi-automated whole-section imaging was performed on a Nikon Eclipse Ni-E epifluorescence microscope using NIS-Elements Advanced Research software (v5.21.03). Images were captured with a Hamamatsu Orca-Fusion Digital Camera (C14440) with a 1×1 binning configuration at 20× magnification (Plan Apo λ, NA = 0.75). Illumination was provided by a Lumencor high color rendering LED light source. For multi-channel fluorescence, a Multilaser Laser Induced Detection Area (LIDA) system was employed. Whole tissue imaging was performed using an automated acquisition pipeline (JOBS) created within NIS-Elements AR (v5.21.03). 4 z-planes, spaced 1.5µm apart, were acquired to capture the optimal focus. All images were saved in Nikon Digital version 2 (ND2) format, which supported large multi-dimensional datasets for downstream analysis.

#### Image Processing

The acquired multi-channel, multi-point, z-stack images were processed with NIS-Elements AR’s (v.5.42.06) Extended Depth of Field (EDF) algorithm to find the best focal point per pixel. The resulting single-plane images were then corrected for uneven illumination and camera artifacts using BaSiC[27] (v1.2.0) and stitched using Ashlar[28] (v1.18.0) based on DAPI channel with *filter-sigma* set to 4 and *maximum-shift* set to 100. Transformation matrices were calculated by detecting and matching local features on DAPI channels of each cycle using AKAZE^22^. All the DAPI channels, including the morphology output of Xenium experiments, were registered to the first cycle of 4i. Transformation matrices were applied using scikit-image’s (v0.26) transform.warp function with linear interpolation.

Throughout this manuscript, we focused on grey matter to reduce technical and biological variability. We manually annotated grey matter boundaries on registered 4i images in QuPath (v0.5.1) using the polygon annotation tool.

#### Cell type annotation

##### Annotating the machine learning dataset

We used deep learning to classify cell types of nuclei from the 4i images. We classified nuclei into 6 different categories: Neurons, Astrocytes, Microglia and Macrophages, Oligodendrocytes and OPCs, cells of vasculature and Unknown. The Uknown were a set of nuclei that did not express any of the relevant features and were intended to be used as a garbage class. To generate the labeled training and validation datasets, we extracted images with the following channels: DAPI (for vasculature), MAP2 (for neurons), CNP (for oligodendrocytes), GFAP (for astrocytes), and Iba1 (for microglia). Additionally, to facilitate annotation of astrocytes, we also overlaid AQP4 and GFAP transcripts on immunofluorescence. These custom extracted channels were annotated into different classes by human annotators by creating point annotations in QuPath (v0.5.1). We found this to be very efficient, since the pixel coordinates of the annotated points can be easily mapped back to Xenium’s segmentation masks. QuPath’s viewer facilitates visualization of the entire tissue which allows the user to sample nuclei from different cortical layers. We annotated 15 different tissue sections, from 15 different individuals, and used annotations from 11 tissue sections for training, 2 for validation and 2 for testing.

##### Generating the training and validation dataset

We reconstructed the Xenium masks with no nuclear expansion as numpy arrays, contrast adjusted the immunofluorescence channels and calculated bounding boxes of each of the labeled masks using regionprops from skimage.measure. We then extracted a 64x64 pixel region, roughly corresponding to 14x14 µm, around the center of the bounding box of each mask for the following channels: DAPI, MAP2, CNP, GFAP and Iba1. From each cropped multi-channel image, we derived two additional spatially informed features: a nucleus-centered proximity map (ring), for providing explicit information about where signals occur relative to the nucleus (i.e. attention), and a locally weighted CNP signal (CNP_local), to resolve confusions caused by close proximity of oligodendrocytes to other nuclei types. These features were derived by exponential distance weighing of the binary nucleus mask, for ring, or CNP channel, for CNP_local.

##### Training the deep learning model

Prior to model input, all multi-channel crops were resized to 224 × 224 pixels to match the spatial requirements of the ResNet architecture. Data augmentation was applied to the training set only and included random horizontal flips, vertical flips, and rotations of up to ±20 degrees to improve model robustness to orientation variability. All channels were normalized independently using a fixed mean and standard deviation of 0.5, effectively scaling intensities to a standardized range. Validation images underwent resizing and normalization only, without augmentation.

A ResNet-18 convolutional neural network pretrained on ImageNet was used as the backbone architecture. To accommodate multi-channel input images, the first convolutional layer was modified to accept 7-channel input while preserving all other kernel parameters. The final layer was modified to correspond to the six cell-type classes. All other network parameters were initialized from pretrained weights. All deep learning steps were performed on NVIDIA A10G on Amazon Web Services (AWS).

We used a weighted cross-entropy loss to address class imbalance. Class weights were computed as the inverse square root of the number of samples per class and normalized to have unit mean. To reduce the influence of the heterogeneous unknown class, its loss contribution was further downweighed by a factor of 0.2. This weighting scheme encouraged balanced learning across cell types while limiting overfitting to poorly defined categories. Learning rate was scheduled using Adam optimizer with an initial rate of 10-4. The learning rate was reduced by half when the validation loss failed to improve for two consecutive epochs. The model was trained for 21 epochs (Supplementary Figure 5b). Performance of the model was assessed using per-class F1-scores.

##### Inference

After training, the model was applied to unlabeled image crops from all the grey matter nuclei in each xenium section, including the subset of nuclei that were used to train the model. The crops were generated in the same way as the training dataset and processed in batches of 1420. Network produced raw class scores were converted to class probabilities using the softmax function, and the class with the highest probability was assigned as the final label. Nuclei with the highest softmax probability below 0.7 were assigned to the Unknown class.

### Spatial transcriptomics data processing and integration

#### Preprocessing of Xenium data

Xenium spatial transcriptomics datasets were processed and analyzed using Seurat (v4.5.2) in R. Xenium output files were loaded using the LoadXenium function and cell type predictions from the above classifier were incorporated into the data. Nuclei with fewer than 20 detected transcripts were excluded from further analysis. Only nuclei found in grey matter were used in downstream analyses.

#### Dimensionality reduction and data visualization

To account for technical and sample-specific variation, batch correction was performed using Harmony, with sample identity specified as the grouping variable. Harmony was applied to the principal component analysis (PCA) embedding, generating a batch-corrected low-dimensional representation. Neighborhood graphs were constructed using Harmony embeddings. Uniform Manifold Approximation and Projection (UMAP) was then computed based on the Harmony reduction to visualize the integrated dataset.

#### Comparison of transcript assignments based on manual cell masks

To determine whether the use of manually annotated, accurate cell masks result in better transcript assignment compared to various methods based on nuclei segmentation in microglia, oligodendrocytes, astrocytes or neurons; we annotated cell boundaries as determined from Iba1, CNP, GFAP and MAP2 immunofluorescence, respectively. Within 3 tissue sections, we annotated 40-41 microglia, 18-25 oligodendrocytes, 15-30 astrocytes, and 40-45 neurons per section from grey matter. We assessed the expression patterns of 32 microglial, 7 oligodendrocyte, 6 astrocytic, 19 neuronal, and 4 vasculature-specific genes within 5µm expansion of the bounding box of manual annotation. Microglial genes were *TMEM119, P2RY12, CX3CR1, TREM2, TYROBP, APOE, GPNMB, SPP1, LPL, CTSS, ABCA7, MS4A4A, MS4A6A, INPP5D, AIF1, CD68, MRC1, ITGAM, C1QA, C1QB, C1QC, C3, HLA-DRA, HLA-DRB1, HLA-DPB1, HLA-DQA1, HLA-DQB1, CD74, CIITA, TNF* and *CXCR4.* Neuronal genes were *RBFOX3, SNAP25, SYT2, SLC17A7, SLC17A6, GAD1, GAD2, MAPT, PVALB, SST, VIP, NPY, RELN, HTR2C, HTR3C, NPAS4, LHX6, GRIK3* and *DRD1*. Oligodendrocyte genes were *MBP, MOG, MOBP, MAG, CNP, OLIG1* and *OLIG2*. Astrocyte genes were *AQP4, ALDH1L1, GLUL, CHI3L1, SERPINA3* and *S100B*. Vasculature-specific genes were *CLDN5, FLT1, PDGFRB* and *RGS5*. True positives, true negatives, false positives, and false negatives were determined based on whether the cell-type specific genes overlap with their respective masks. F1-scores were averaged between samples.

## ETHICS APPROVAL AND CONSENT TO PARTICIPATE

This study was approved by the Institutional Review Board of Columbia University (AAAR4962). All human tissue donors provided informed consent, permitting the use of their postmortem brains and associated antemortem clinical data for research purposes.

## AUTHOR CONTRIBUTIONS

PLD, VM, HK, MT, and YZ conceived and designed the study. BK designed the 4i panel and the Nikon JOBS pipeline. TL performed the 4i experiments and image acquisition. WC and TL prepared samples for the Xenium experiment under the supervision of YZ. BK and AW performed sample registration and alignment for 4i (post-Xenium) and Xenium datasets. BK, AW, and HK analyzed the data. DB provided the samples. BK, TL, and MT drafted the manuscript. All authors interpreted the results and approved the final version of the manuscript.

## FUNDING INFORMATION

The study was supported by NIH grant U19 AG074862. ROSMAP is supported by P30AG10161, P30AG72975, R01AG17917, R01AG015819, U01AG072572, and U01AG046152.

## ACKNOWLEDGEMENTS

We thank the participants in the ROS and MAP studies for their generous contribution to the study of cognitive aging and AD.

## COMPETING INTERESTS

The authors declare no conflicts of interest.

## Notes

### Competing Interest Statement

The authors have declared no competing interest.

