## Supplementary Figures and Tables for "High-Dimensional Multi-Omic Mapping of Post-Mortem Human Brain Using Iterative Indirect Immunofluorescence Imaging on Xenium-Processed Tissues"

Supplementary Figure 1

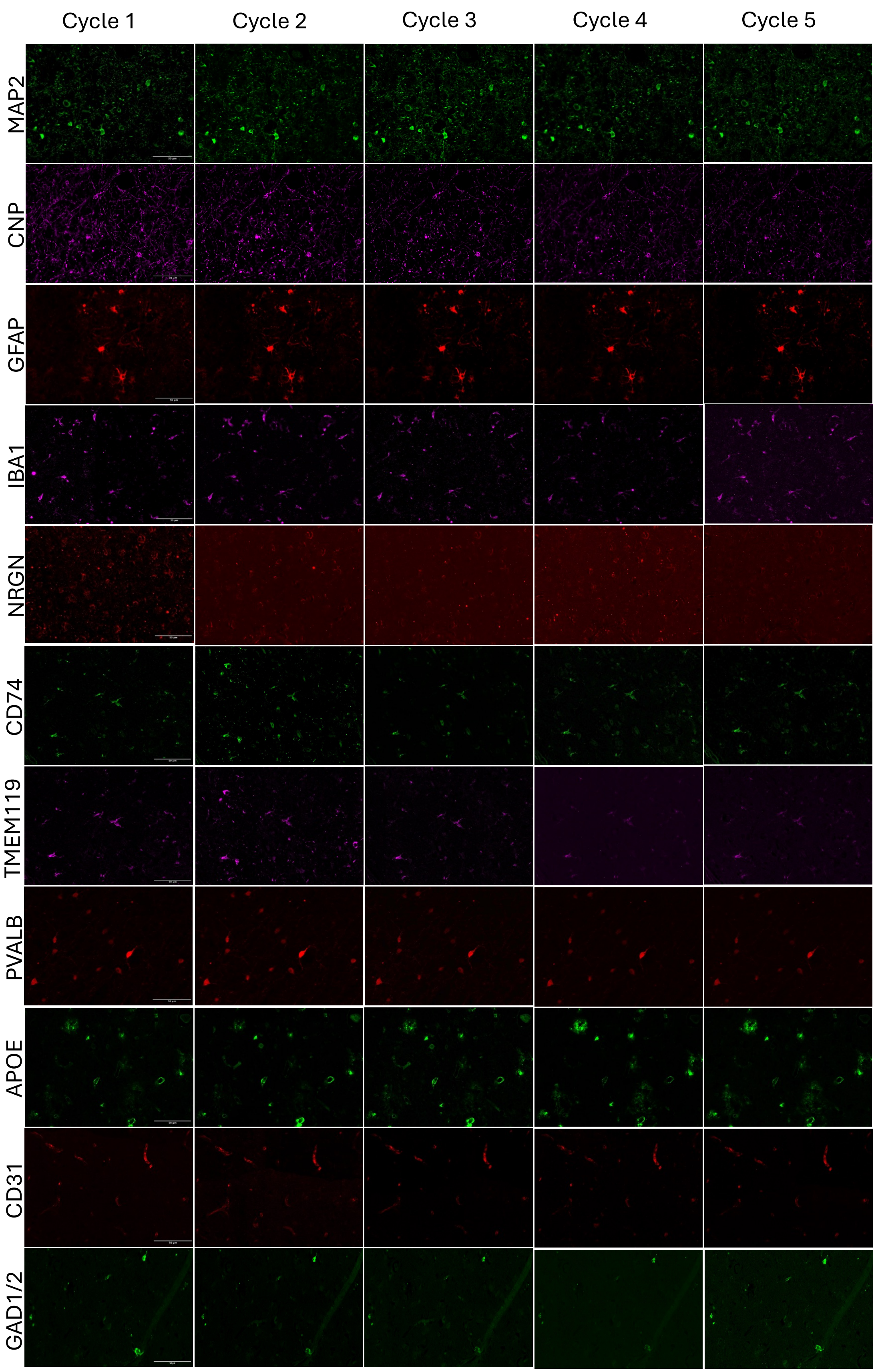

Supplementary Figure 2

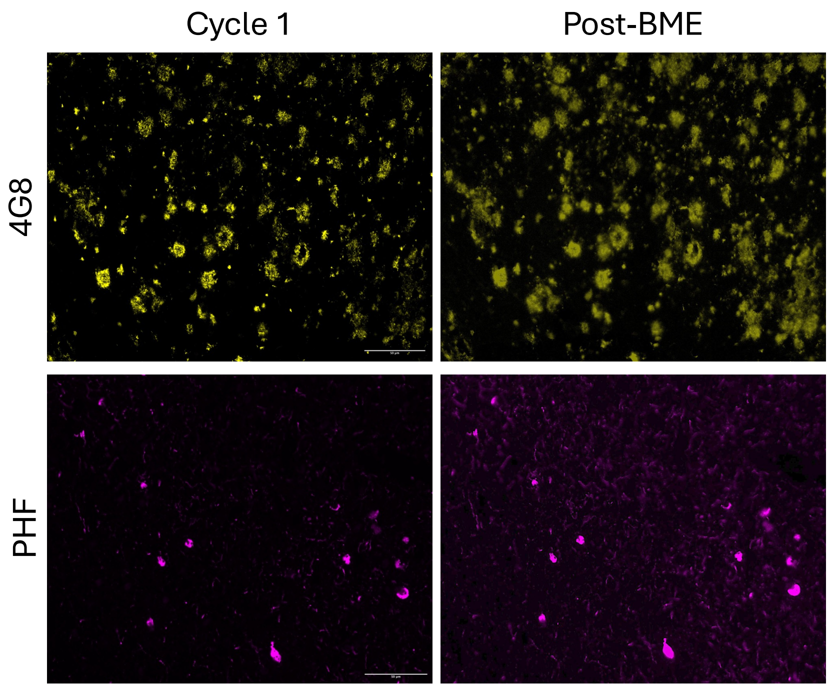

Supplementary Figure 3

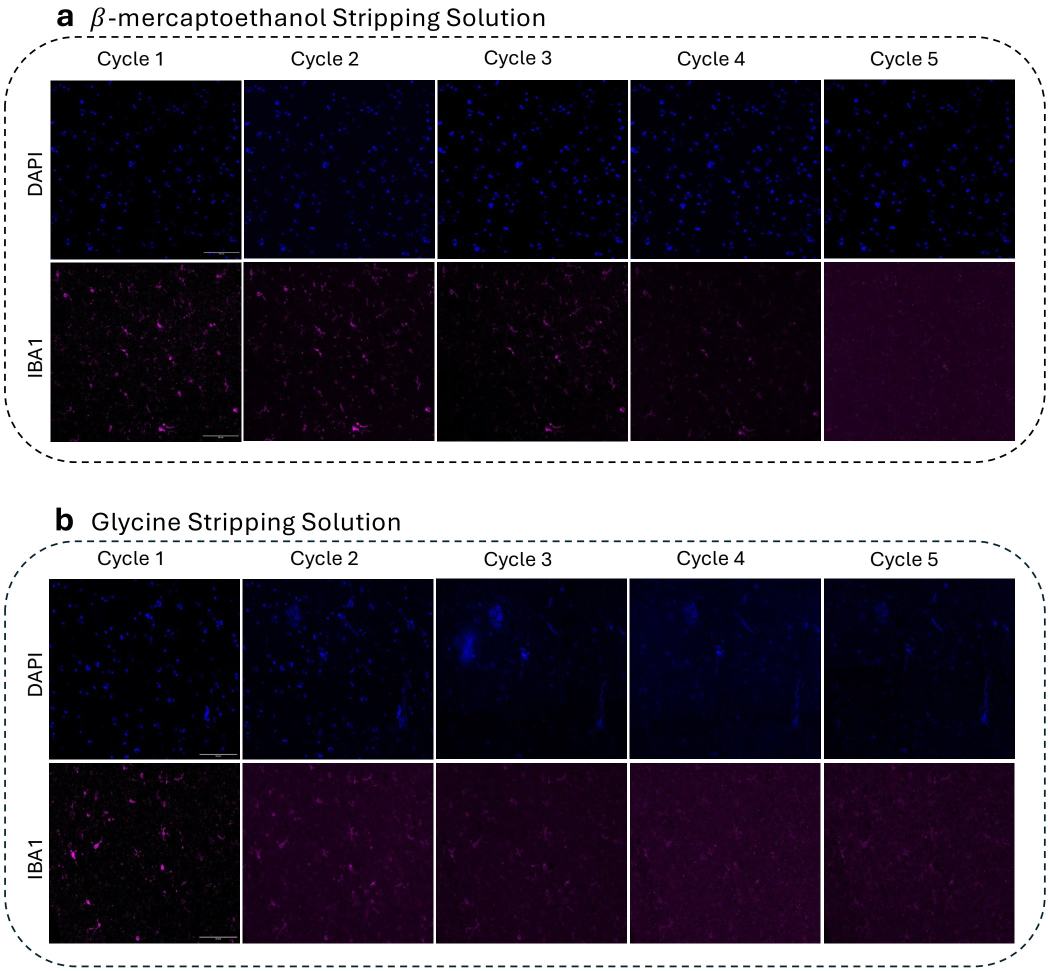

### Supplementary Figure 4

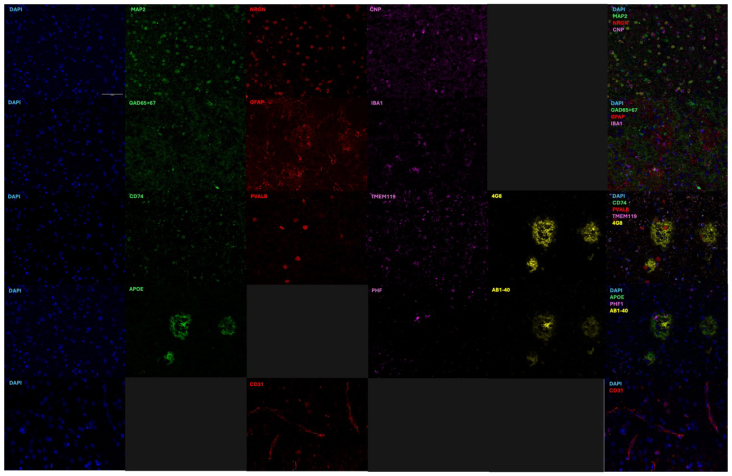


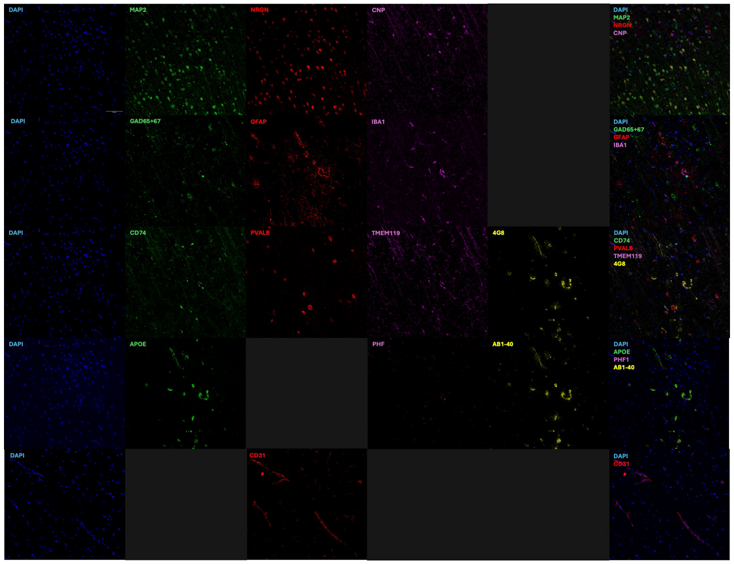

Supplementary Figure 5

**a**

| | Marker | Training n = 11 | Validation n = 2 | Testing n = 2 |
|---|---|---|---|---|
| Neurons | MAP2 | 35652 | 6682 | 6352 |
| Microglia | Iba1 | 6840 | 1255 | 1039 |
| Astrocytes | GFAP | 5447 | 633 | 1331 |
| Oligodendrocytes | CNP | 5493 | 1157 | 1062 |
| Vasculature | Morphology | 4041 | 888 | 501 |
| Unknown | N/A | 1626 | 294 | 207 |
| **Total** | | **59,099** | **10,909** | **10,492** |

**b**

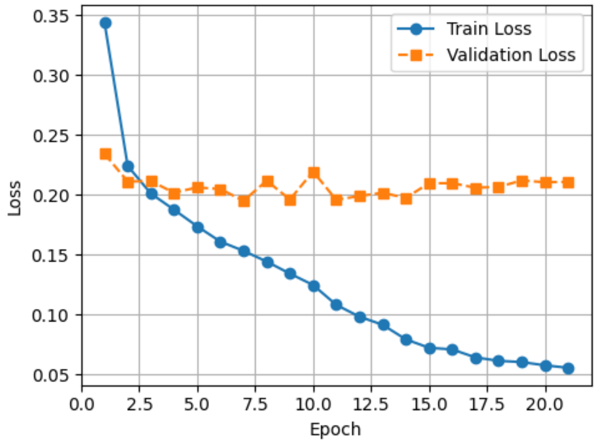


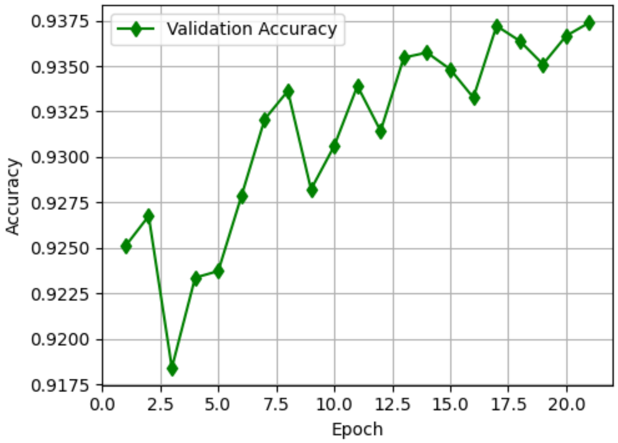

Supplementary Figure 6

a

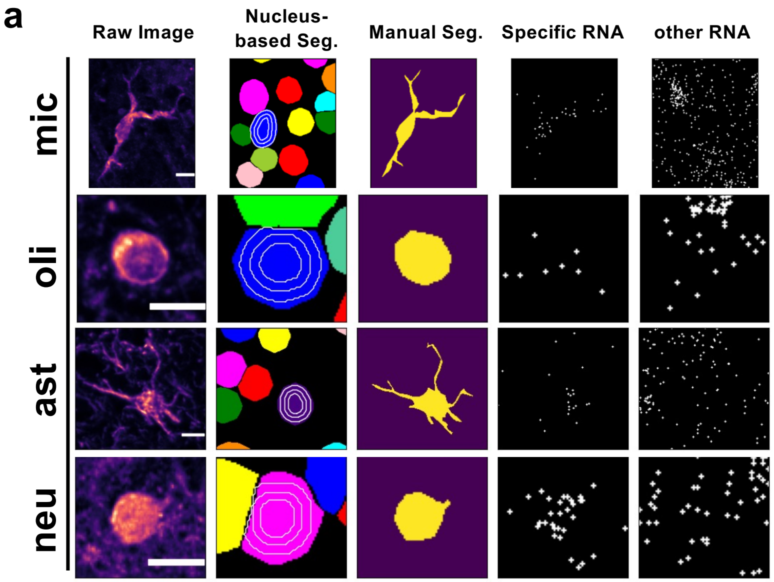


b

| | Manual Seg. | Nucleus Only | 5µm Expansion | 50% Expansion | 100% Expansion |
|---|---|---|---|---|---|
| **f1** | | | | | |
| mic | 0.525 | 0.304 | 0.535 | 0.459 | 0.508 |
| oli | 0.473 | 0.450 | 0.423 | 0.454 | 0.436 |
| ast | 0.304 | 0.142 | 0.327 | 0.241 | 0.300 |
| neu | 0.686 | 0.386 | 0.769 | 0.684 | 0.763 |
| **nCount** | | | | | |
| mic | 16.675 | 6.525 | 18.877 | 12.581 | 16.315 |
| oli | 5.199 | 2.652 | 8.402 | 5.086 | 6.741 |
| ast | 14.180 | 4.482 | 14.041 | 8.696 | 12.821 |
| neu | 28.694 | 12.669 | 36.238 | 27.786 | 35.181 |

### Supplementary Table 1

| projid | ID | Age of Death | Sex | NIA-Reagan | Cognition | CAA | APOE genotype |
|---|---|---|---|---|---|---|---|
| 02899847 | xenium_1 | 74 | male | Low | MCI | Moderate | 33 |
| 03713990 | xenium_2 | 88 | male | Int. | NCI | Mild | 33 |
| 10101741 | xenium_3 | 90+ | male | High | AD | Mild | 33 |
| 10248033 | xenium_4 | 88 | male | Int. | AD | Moderate | 44 |
| 10425642 | xenium_5 | 87 | male | High | AD | Mild | 33 |
| 17929065 | xenium_6 | 90+ | female | Low | NCI | Mild | 33 |
| 18414513 | xenium_7 | 90+ | male | Int. | AD | Moderate | 33 |
| 20372539 | xenium_8 | 86 | female | High | AD | Severe | 44 |
| 33321607 | xenium_9 | 90+ | male | Low | NCI | Mild | 33 |
| 50302392 | xenium_10 | 90+ | female | High | AD | Mild | 33 |
| 50401002 | xenium_11 | 86 | female | Int. | NCI | Mild | 33 |
| 54122640 | xenium_12 | 90+ | female | High | AD | Moderate | 33 |
| 59150662 | xenium_13 | 90+ | female | Int. | AD | Mild | 33 |
| 77180612 | xenium_14 | 87 | female | High | AD | Severe | 33 |
| 94328246 | xenium_15 | 90+ | male | High | MCI | Moderate | 33 |

Supplementary Table 2

| Cycle | Fluorophore | | | |
|---|---|---|---|---|
| | **Alexa Fluor 488** | **Alexa Fluor 568** | **Alexa Fluor 647** | **Alexa Fluor 790** |
| **1** | MAP2 | NRGN | CNP | |
| **2** | GAD65+GAD67 | GFAP | IBA1 | |
| **3** | CD74 | PVALB | TMEM119 | β-Amyloid, 17-24 |
| **4** | APOE | | Tau (phospho S396) | β-Amyloid, 1-40 |
| **5** | | CD31 | | |

### Supplementary Table 3

| REAGENT or RESOURCE | SOURCE | IDENTIFIER |
|---|---|---|
| Antibodies | | |
| Chicken polyclonal anti-MAP2 (1:500) | Novus Biologicals | Cat#NB300-213; RRID: AB_2138178 |
| Mouse monoclonal anti-NRGN [C2] (1:100) | Invitrogen | Cat#MA5-41648; RRID: AB_2899130 |
| Rabbit polyclonal anti-CNP (1:100) | Sigma-Aldrich | Cat#HPA023278; RRID: AB_1847060 |
| Rabbit monoclonal anti-GAD65+GAD67 [EPR19366] (1:100) | Abcam | Cat#ab183999; RRID: AB_3662875 |
| Mouse monoclonal anti-GFAP, Cy3 conjugate (1:100) | Sigma-Aldrich | Cat#MAB3402C3; RRID: AB_11213580 |
| Goat polyclonal anti-IBA1 (1:100) | FUJIFILM Wako Pure Chemical Corporation | Cat#011-27991; RRID: AB_2935833 |
| Rabbit polyclonal anti-CD74 (1:100) | Sigma-Aldrich | Cat#HPA010592; RRID: AB_1078482 |
| Sheep polyclonal anti-PVALB (1:300) | Invitrogen | Cat#PA5-47693; RRID: AB_2609239 |
| Rabbit monoclonal Alexa Fluor® 647 anti-TMEM119 [EPR25865-89] (1:100) | Abcam | Cat#ab313674 |
| Mouse monoclonal anti-β-Amyloid, 17-24 (1:16000) | BioLegend | Cat#800702; RRID: AB_2564634 |
| Goat polyclonal anti-APOE (1:300) | Invitrogen | Cat#PA5-18361; RRID: AB_10979861 |
| Rabbit monoclonal anti-Tau (phospho S396) [EPR2731] (1:100) | Abcam | Cat#ab156623 |
| Mouse monoclonal anti-β-Amyloid, 1-40 (1:300) | BioLegend | Cat#805402; RRID: AB_2564681 |
| Rabbit polyclonal anti-CD31 (1:50) | Abcam | Cat#ab28364; RRID: AB_726362 |
| Donkey anti-Rabbit IgG (H+L) Highly Cross-Adsorbed Secondary Antibody, Alexa Fluor™ 488 (1:500) | Invitrogen | Cat#A-21206; RRID: AB_2535792 |
| Donkey anti-Goat IgG (H+L) Cross-Adsorbed Secondary Antibody, Alexa Fluor™ 488 (1:500) | Invitrogen | Cat#A-11055; RRID: AB_2534102 |
| Donkey anti-Chicken IgY (H+L) Highly Cross Adsorbed Secondary Antibody, Alexa Fluor™ 488 (1:500) | Invitrogen | Cat#A78948; RRID: AB_2921070 |
| Donkey anti-Mouse IgG (H+L) Highly Cross-Adsorbed Secondary Antibody, Alexa Fluor™ 568 (1:500) | Invitrogen | Cat#A10037; RRID: AB_11180865 |
| Donkey anti-Rabbit IgG (H+L) Highly Cross-Adsorbed Secondary Antibody, Alexa Fluor™ 568 (1:500) | Invitrogen | Cat#A10042; RRID: AB_2534017 |
| Donkey anti-Sheep IgG (H+L) Cross-Adsorbed Secondary Antibody, Alexa Fluor™ 568 (1:500) | Invitrogen | Cat#A-21099; RRID: AB_2535753 |
| Donkey anti-Rabbit IgG (H+L) Highly Cross-Adsorbed Secondary Antibody, Alexa Fluor™ 647 (1:500) | Invitrogen | Cat#A-31573; RRID: AB_2536183 |
